# VCFcontam: A Machine Learning Approach to Estimate Cross-Sample Contamination from Variant Call Data

**DOI:** 10.1101/2021.03.12.435007

**Authors:** Evan McCartney-Melstad, Ke Bi, James Han, Catherine K. Foo

## Abstract

The quality of genotyping calls resulting from DNA sequencing is reliant on high quality starting genetic material. One factor that can reduce sample quality and lead to misleading genotyping results is genetic contamination of a sample by another source, such as cells or DNA from another sample of the same or different species. Cross-sample contamination by individuals of the same species is particularly difficult to detect in DNA sequencing data, because the contaminating sequence reads look very similar to those of the intended base sample. We introduce a new method that uses a support vector regression model trained on *in silico* contaminated datasets to predict empirical contamination using a collection of variables drawn from VCF files, including the fraction of sites that are heterozygous, the fraction of heterozygous sites with imbalanced allele counts, and parameters describing distributions fit to heterozygous allele fractions in a sample. We use the method described here to train a model that can accurately predict the extent of cross-sample contamination within 1% of the actual fraction, for simulated contaminated samples in the 0-5% contamination range, directly from the VCF file.

**Definitions:** *Lesser allele:* The allele in a heterozygous position that received less sequencing read support (which may be either the REF or ALT allele).

*Lesser allele fraction (LAF):* The number of sequencing reads supporting the less frequently observed allele divided by the sum of reads supporting both alleles in the genotype at a given genomic position.

## Introduction

Avoiding cross-sample contamination is critical for generating accurate and high-quality next-generation sequencing (NGS) data. The presence of contaminating DNA from the same or closely related species can lead to clinical false positives (Mendez et al., 2019). However, it is often impossible to tell if a sample has been contaminated in the laboratory, and so the raw data collected for a sample must itself be analyzed to look for evidence of contamination.

In a diploid organism, three genotypes are possible for autosomal genomic locations: homozygous for the reference (REF) allele, heterozygous, or homozygous for an alternative (ALT) allele. For sites where a diploid individual is homozygous for the REF allele, the ideal expectation is that all sequenced bases at the site should represent the REF allele, and the same is expected for observed ALT bases for homozygous ALT sites. Because paternally and maternally inherited autosomes are present at an even ratio in cells, for heterozygous sites the ideal expectation is that 50% of the data generated for a heterozygous site should represent one allele and the other 50% should represent the other allele. Some deviation from this pattern is generally expected, particularly for heterozygous sites, due to factors including sequencing error and sampling error.

Sequencing a sample contaminated by DNA from another sample, however, will lead to more significant deviations from the expected 50:50 ratio in heterozygotes and can lead to the observation of ALT read support in homozygous REF sites and REF read support in homozygous ALT sites. The degree to which contamination affects these observations depends on the fraction of chromosome copies in a DNA sample that arise from the base sample versus the contaminating sample. Here we demonstrate a new method that uses the information present in these biased allele fractions to estimate continuous numerical cross-sample contamination fractions.

Two recent methods have been published that also perform continuous estimations of cross-sample allele fractions. VerifyBamID2 (Zhang et al., 2020) co-estimates genetic ancestry and contamination fraction from BAM files. ART-DeCo is another promising tool that offers a non-parametric worst-case contamination fraction estimate based on lesser allele fractions, although it reportedly has low specificity and requires possible contaminant context in order to confirm positive contamination (Fiévet et al., 2019). A third tool that also uses machine learning based on modelling lesser allele fractions has been released in the software package *vanquish* (Jiang et al. 2019), although this tool classifies samples as contaminated or not contaminated, rather than estimating a continuous contamination fraction. Our method is distinct from these tools in that it estimates a continuous contamination fraction from VCF files with high accuracy even at low contamination fractions.

## Methods

### Estimation method

We estimate the contamination fraction by modelling heterozygosity and deviations from the expected allelic ratios across a curated list of highly polymorphic genomic locations. The list of genomic locations consists of single nucleotide polymorphisms (SNP) that are likely to differ between any two sampled alleles. While any list of highly polymorphic loci could be used for this purpose, as a demonstration, we limit our analysis here to chromosome 21 and select the 23,234 genomic locations that meet the following characteristics based on the genome VCF data from gnomAD v2.1.1 (Karczewski et al., 2020):

1. Biallelic SNP, to exclude sites that have multiallelic or insertion/deletion variation
2. Filter == PASS
3. Minor allele frequency >= 35%, to select for highly polymorphic loci

To analyze a sample for contamination, we begin with a VCF file in which all positions (including homozygous reference) have been emitted by the variant caller. Homozygous reference sites may contain information relevant to contamination, and so we use all-sites VCFs here. (If homozygous reference sites are absent from the VCFs, this method could be revised to omit this variable from the model.) For each of the previously curated positions in a given all-sites VCF, we then calculate the allelic depths supporting both the REF and ALT alleles, recording the following:

1. *heterozygosity*: The fraction of sites that are heterozygous
2. *fraction_heterozygous_minABfracBelow20*: The number of heterozygous calls with lesser allele fraction (LAF) < 20%, divided by the total number of heterozygous calls
3. *fraction_homozygous_minAB0*: The number of homozygous REF sites with zero ALT allele counts, divided by the total number of homozygous REF calls
4. A list of lesser allele fractions for all sites that are either a) heterozygous, or b) homozygous with non-zero lesser allele counts

A lognormal mixture model is then fit to the list of lesser allele fractions (#4 described above) using the functions `GeneralMixtureModel()` and `fit()` from the python library `pomegranate` (Schreiber, 2017). Because true diploid heterozygotes are expected to result in lesser allele frequencies close to 50%, and contamination is expected to result in lesser allele frequencies closer to 0%, a bimodal distribution of lesser allele frequencies is expected in the presence of appreciable contamination. To quantify this, a mixture model is first initialized with two lognormal distributions to attempt to capture a bimodal distribution if present, the first with a mean of 0 and a standard deviation of 1, and the second with a mean of 0.5 and a standard deviation of 1. The mixture model is then fitted to the observed lesser allele fractions. After fitting, the means and standard deviations of the lognormal distributions are recorded, along with their respective weights from the mixture model.

A support vector regression model (Drucker et al., 1997) is then trained using a large collection of *in silico* contaminated samples (described below) using the `svm.SVR()` function from `scikit-learn` (Pedregosa et al., 2011). The response variable is the percentage of reads in the sample that are from a contaminating sample. The predictor variables include:

1. *heterozygosity*
2. *fraction_heterozygous_minABfracBelow20*
3. *fraction_homozygous_minAB0*
4. The means (*mu1*, *mu2*) and weights (*weights1*, *weights2*) of the two distributions in the lognormal mixture model as described above

After training the support vector regression modeling using *in silico* contaminated samples, the model is serialized as a python object. Then, for any new sample VCF, the predictor variables described above are calculated by parsing the VCF file with pysam (https://github.com/pysam-developers/pysam) as done when generating the training data. The values calculated for the VCF file are then input into the serialized support vector regression model and the contamination fraction is estimated.

### in silico *contaminated dataset for training and testing the estimation model*

To train and test the estimation model, we created a dataset of *in silico* contaminated VCFs. These VCFs were derived from the 1000 Genomes dataset sequenced by the New York Genome Center (30x whole genome sequencing, PRJEB31736). A collection of 51 CRAM (Hsi-Yang Fritz et al., 2011) files from different individuals (Appendix A1) were downloaded, subset to read pairs mapping to chromosome 21, and converted to 150bp paired-end FASTQ files using samtools v1.10 (Li et al., 2009).

From these 51 starting samples, a set of 500 *in silico* contaminated R1/R2 FASTQ files were generated by combining reads from two randomly drawn samples at specific ratios. The average coverage and the extent of contamination in the training and testing datasets should be designed to match the characteristics of the data for which the user intends to utilize the estimator; for our purposes, we chose to focus on contamination percentages of 0-5% and coverage values around 20X. We generated a range of contamination percentages that followed a beta distribution with shape parameters 1 and 5, ranging from 0.036% to 13.816%, and selected total read counts for the *in silico* mixtures from a uniform distribution between 6 and 9 million reads (all mapping to chromosome 21; Figure 1). This set of 500 *in silico* contaminated samples were then randomly partitioned into a training set of 300 FASTQ samples and a testing set of 200 FASTQ samples.

**Figure 1:**
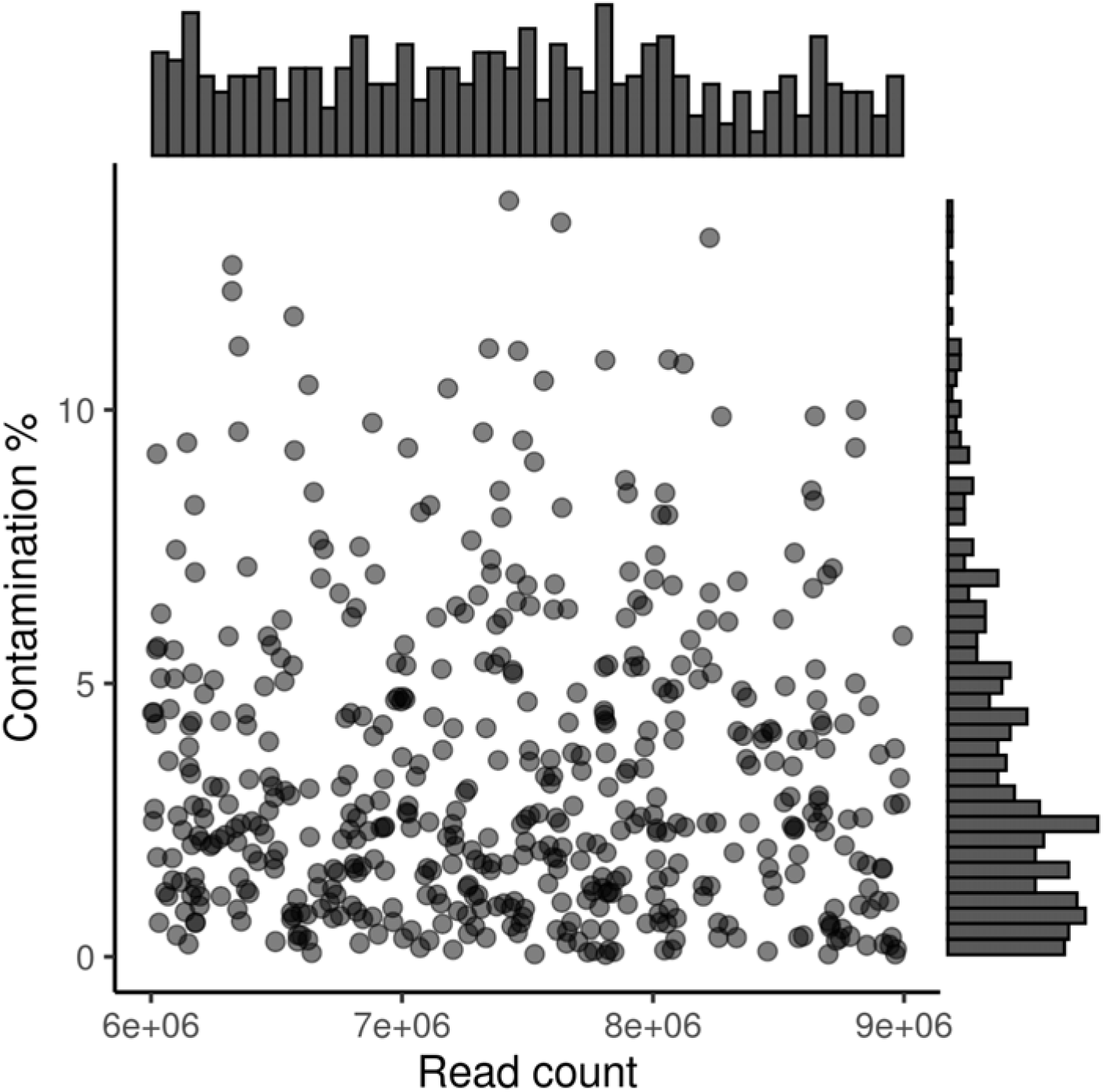
*in silico* simulated dataset characteristics. Read counts are the number of reads in the final contaminated samples. The bar graphs are histograms for both axes.

As expected, lesser allele fractions showed markedly different patterns in highly contaminated samples compared to relatively lightly contaminated ones (Figure 2). Visualizing these minor allele fractions, along with the lognormal distributions fit to them, make it apparent that both the relative heights and the relative positions of the two distributions are likely to be predictive of the amount of cross-sample contamination.

**Figure 2:**
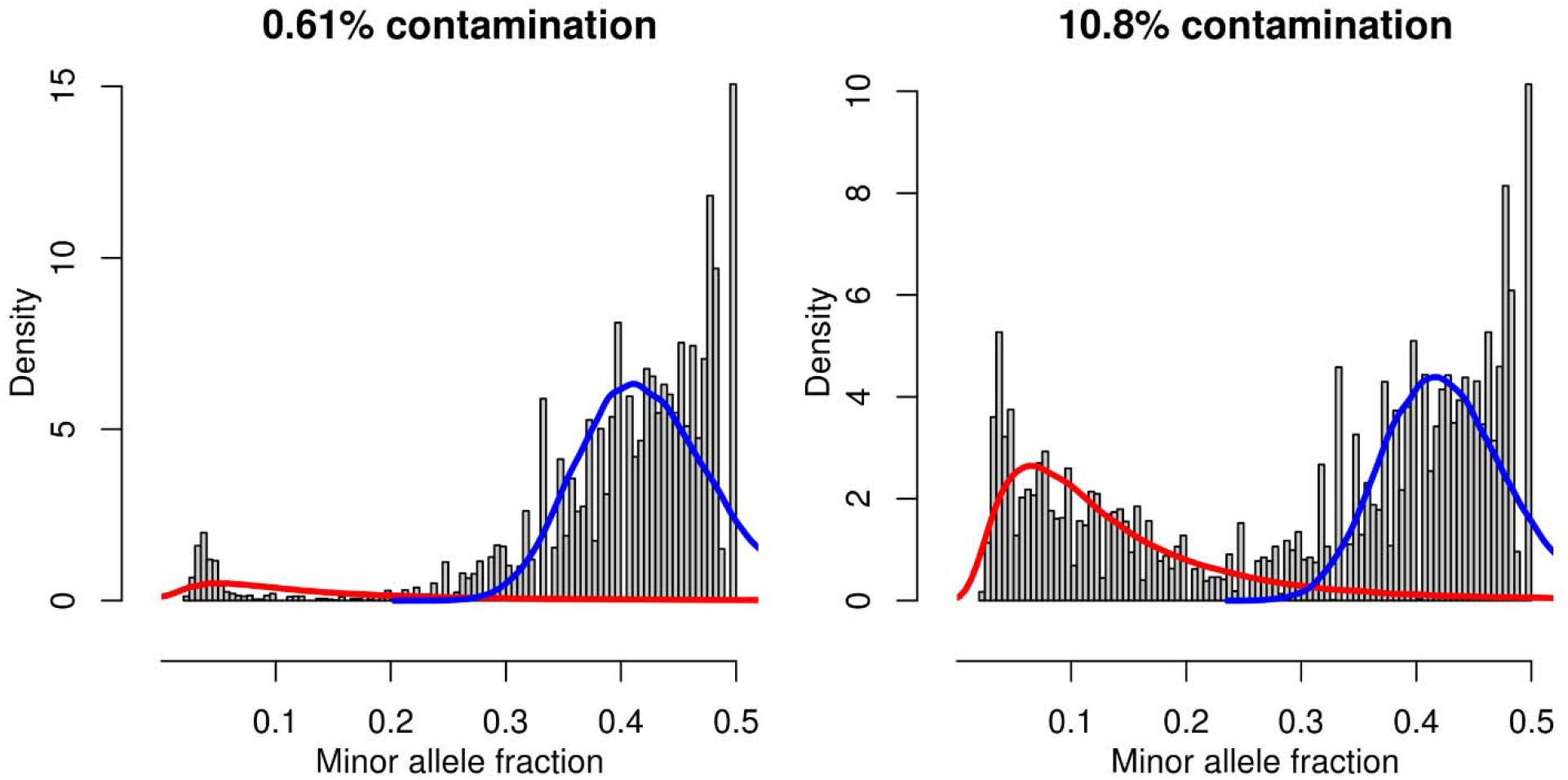
Examples of minor allele distributions for a low contamination (0.61%, at left) and high contamination (10.8%, at right) sample. Histograms represent the observed minor allele fractions in qualifying sites for two different simulated contamination levels. The red and blue lines represent fitted lognormal distributions with y-axis values scaled by the respective weights in the mixture model. The more highly contaminated sample has a noticeably higher fraction of sites with low minor allele fractions.

### Training the model

Each *in silico* contaminated sample was genotyped for the list of SNPs described above, by creating single-sample VCF files using DRAGEN v.3.3.8f (Miller et al., 2019). The positions selected on chromosome 21 were used as the variant calling BED for computational efficiency (although any variant calling region of interest that included the chosen intermediate-frequency sites would be acceptable). All relevant features described in the *Estimation method* section were extracted from each of the 300 training VCFs (Appendix A2). A support vector regression model was then trained on the extracted features and the known simulated contamination values and serialized. Finally, contamination estimates were predicted for the 200 remaining *in silico* contaminated samples in the testing dataset (extracted features for the testing dataset are in Appendix A3).

## Results

Predicted values for the 300 testing VCFs were highly concordant with simulated values (adjusted R^2^ = 0.9528), particularly in the 0% to 8% range (Figure 3 and Appendix A3). The mean absolute difference between the predicted and simulated contamination value was 0.37% (0.28% for the simulated range between 0% and 8%). The maximum prediction error was 1.37% in the 0%-8% range, and 4.31% overall. In total, only 11 out of 200 predictions had an error greater than 1 percentage point, and just 4 of these were within the 0%-8% simulated contamination range. Above 8%, the model tended to underpredict contamination by several percentage points. Run time for the VCFs in the example dataset described here is approximately 15-20 seconds on a single processor core, consuming about 125 MB of RAM.

**Figure 3:**
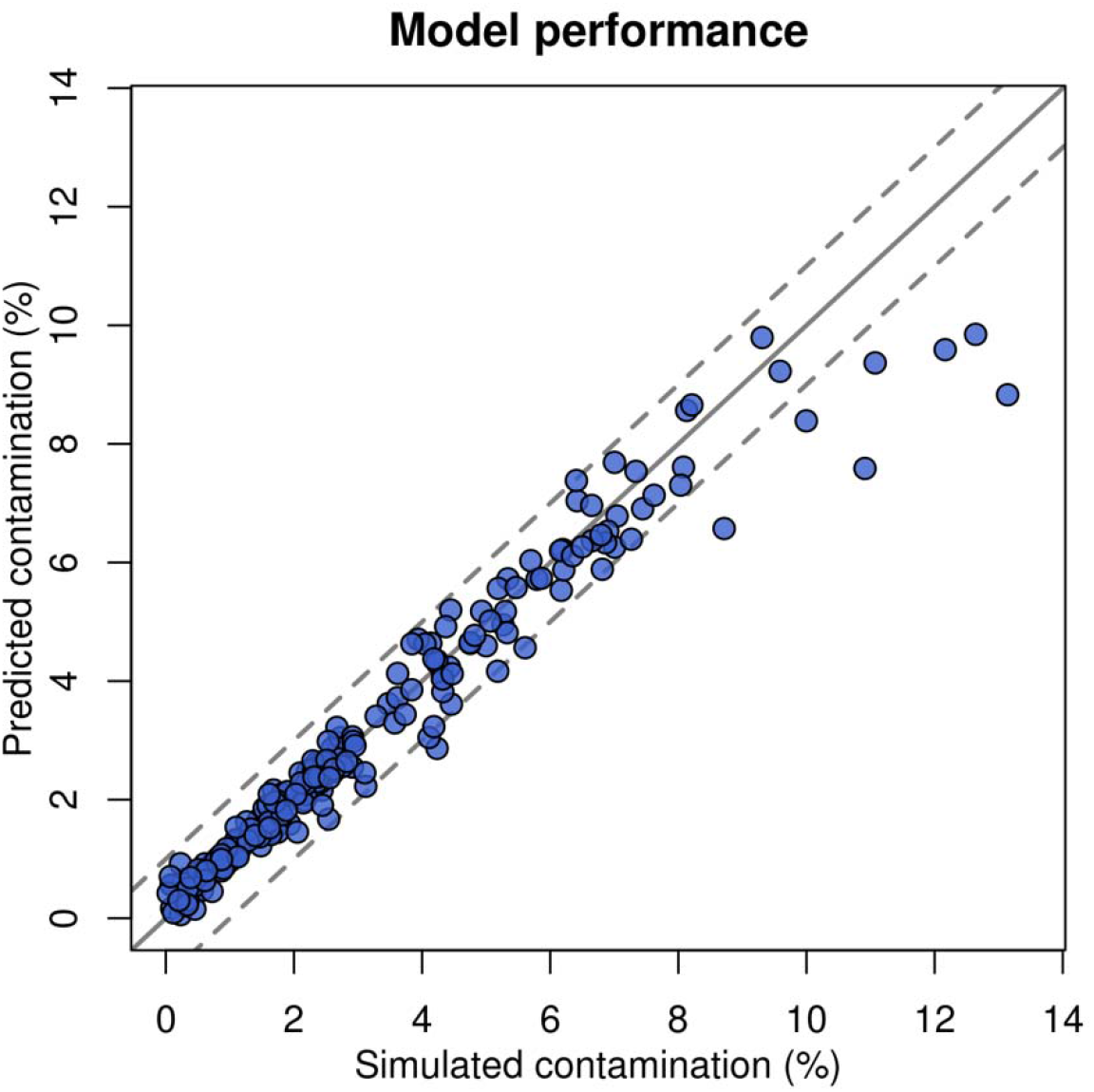
Predicted versus simulated contamination percentages

## Discussion

Cross-sample contamination is a quality metric monitored in many laboratories, in order to ensure that samples with contamination levels that may interfere with variant calling accuracy or other results are flagged and removed from the pipeline prior to downstream biological or clinical interpretation. Here we describe a method that uses lesser allele fractions from a VCF to estimate the level of contamination. This utility allows an accurate evaluation of sample-wide contamination levels without access to the underlying FASTQ or BAM/CRAM data.

This work provides a framework for the creation of a contamination estimator tailored to any particular application. The list of polymorphic sites for which the lesser allele fractions are collected can follow the guidelines described here, or can be altered to include sites of known polymorphism present in the population being studied. The training dataset should be created with contamination and coverage characteristics that match the anticipated characteristics of the samples to be evaluated. In this demonstration, we wanted to develop an estimator that would have high accuracy in 0-5% contamination range, where we had previously observed a decline of performance in variant calling accuracy (data not shown). As a result, we designed our *in silico* dataset to enrich for samples with contamination values in this range (Figure 1). Accuracy for higher contamination values was not as critical for our application, and thus it was not surprising that the accuracy was poor above 8%, given the lack of training data at that contamination level (Figure 3).

The *in silico* contaminated dataset here also focused on a coverage level of about 20X. Since the allelic depths and random sampling bias are sensitive to the level of coverage, further work is needed to determine whether this approach can still result in accurate performance at much lower or much higher levels of coverage.

The method as implemented here assumes that contamination occurs between two genetically unrelated samples. If the source of contamination is the DNA of a close relative, then the contamination levels are likely to be underestimated.

## Supporting information

Appendix A2: Training data features

Appendix A3: Testing data features and estimates

## Software Availability

VCFcontam is available at: https://github.com/Ancestry/VCFcontam

## Acknowledgements

All work was supported and funded by AncestryDNA. DNA sequence data used to generate the *in silico* contamination experiments were generated at the New York Genome Center with funds provided by NHGRI Grant 3UM1HG008901-03S1. Data from NA12046, NA12342, NA12761, NA12815, NA12878, NA12889, and NA12890 were generated by the New York Genome Center from cell lines obtained from the NIGMS Human Genetic Cell Repository at the Coriell Institute for Medical Research.

## Appendix

A1: Individuals included in in silico analysis

- HG00451
- HG00583
- HG00626
- HG00634
- HG00708
- HG00731
- HG00732
- HG00982
- HG01198
- HG01323
- HG01390
- HG01414
- HG01443
- HG01465
- HG01468
- HG01522
- HG01530
- HG01890
- HG01976
- HG02006
- HG02253
- HG02278
- HG02355
- HG02379
- HG02445
- HG02604
- HG02759
- HG02775
- HG03021
- HG03045
- HG03077
- HG03442
- HG03548
- HG03716
- HG03765
- HG03815
- HG03832
- HG03851
- HG03965
- HG03977
- NA12046
- NA12342
- NA12761
- NA12815
- NA12878
- NA12889
- NA12890
- NA18510
- NA18548
- NA18557

A2: Features extracted from the 300 training VCFs

A3: Features extracted from, and estimated contamination for, the 200 testing VCFs

See Supplementary Materials

## References

Drucker, H., Burges, C.J., Kaufman, L., Smola, A., Vapnik, V., 1997. Support vector regression machines. Adv. Neural Inf. Process. Syst. 9, 155–161.

Fiévet, A., Bernard, V., Tenreiro, H., Dehainault, C., Girard, E., Deshaies, V., Hupe, P., Delattre, O., Stern, M.-H., Stoppa-Lyonnet, D., Golmard, L., Houdayer, C., 2019. ART-DeCo: easy tool for detection and characterization of cross-contamination of DNA samples in diagnostic next-generation sequencing analysis. Eur. J. Hum. Genet. 27, 792–800. https://doi.org/10.1038/s41431-018-0317-x

Hsi-Yang Fritz, M., Leinonen, R., Cochrane, G., Birney, E., 2011. Efficient storage of high throughput DNA sequencing data using reference-based compression. Genome Res. 21, 734–740. https://doi.org/10.1101/gr.114819.110

Jiang, T., Buchkovich, M., Motsinger-Reif, A. Same-species contamination detection with variant calling information from next generation sequencing. bioRxiv 531558. https://doi.org/10.1101/531558

Karczewski, K.J., Francioli, L.C., Tiao, G., Cummings, B.B., Alföldi, J., Wang, Q., Collins, R.L., Laricchia, K.M., Ganna, A., Birnbaum, D.P., Gauthier, L.D., Brand, H., Solomonson, M., Watts, N.A., Rhodes, D., Singer-Berk, M., England, E.M., Seaby, E.G., Kosmicki, J.A., Walters, R.K., Tashman, K., Farjoun, Y., Banks, E., Poterba, T., Wang, A., Seed, C., Whiffin, N., Chong, J.X., Samocha, K.E., Pierce-Hoffman, E., Zappala, Z., O’Donnell-Luria, A.H., Minikel, E.V., Weisburd, B., Lek, M., Ware, J.S., Vittal, C., Armean, I.M., Bergelson, L., Cibulskis, K., Connolly, K.M., Covarrubias, M., Donnelly, S., Ferriera, S., Gabriel, S., Gentry, J., Gupta, N., Jeandet, T., Kaplan, D., Llanwarne, C., Munshi, R., Novod, S., Petrillo, N., Roazen, D., Ruano-Rubio, V., Saltzman, A., Schleicher, M., Soto, J., Tibbetts, K., Tolonen, C., Wade, G., Talkowski, M.E., Neale, B.M., Daly, M.J., MacArthur, D.G., 2020. The mutational constraint spectrum quantified from variation in 141,456 humans. Nature 581, 434–443. https://doi.org/10.1038/s41586-020-2308-7

Li, H., Handsaker, B., Wysoker, A., Fennell, T., Ruan, J., Homer, N., Marth, G., Abecasis, G., Durbin, R., 1000 Genome Project Data Processing Subgroup, 2009. The Sequence Alignment/Map format and SAMtools. Bioinformatics 25, 2078–2079. https://doi.org/10.1093/bioinformatics/btp352

Mendez, F.L., Jiang, R., White, S., Lee, W., 2019. BACON: Baited Abrogation of CONtamination. Poster at American Society of Human Genetics Meeting, Houston, TX.

Miller, N.A., Farrow, E.G., Gibson, M., Willig, L.K., Twist, G., Yoo, B., Marrs, T., Corder, S., Krivohlavek, L., Walter, A., Petrikin, J.E., Saunders, C.J., Thiffault, I., Soden, S.E., Smith, L.D., Dinwiddie, D.L., Herd, S., Cakici, J.A., Catreux, S., Ruehle, M., Kingsmore, S.F., 2015. A 26-hour system of highly sensitive whole genome sequencing for emergency management of genetic diseases. Genome Med. 7, 100. https://doi.org/10.1186/s13073-015-0221-8

Pedregosa, F., Varoquaux, G., Gramfort, A., Michel, V., Thirion, B., Grisel, O., Blondel, M., Prettenhofer, P., Weiss, R., Dubourg, V., Vanderplas, J., Passos, A., Cournapeau, D., Brucher, M., Perrot, M., Duchesnay, É., 2011. Scikit-learn: Machine Learning in Python. J. Mach. Learn. Res. 12, 2825–2830.

Schreiber, J., 2017. Pomegranate: fast and flexible probabilistic modeling in python. J. Mach. Learn. Res. 18, 5992–5997.

Zhang, F., Flickinger, M., Taliun, S.A.G., Consortium, I.P.G., Abecasis, G.R., Scott, L.J., McCaroll, S.A., Pato, C.N., Boehnke, M., Kang, H.M., 2020. Ancestry-agnostic estimation of DNA sample contamination from sequence reads. Genome Res. https://doi.org/10.1101/gr.246934.118

